# Conformalized Multiview Learning

**DOI:** 10.64898/2026.01.22.700165

**Authors:** Saptarshi Roy, Sreya Sarkar, Chitrak Banerjee, Erina Paul, Piyali Basak, Himel Mallick

## Abstract

There is an overwhelmingly large body of literature and numerous algorithms already available on “multimodal AI”, based on different modeling and fusion paradigms. As multimodal AI becomes fully integrated into healthcare applications, the ability to say “I’m not sure” or “I don’t know” when uncertain is a necessary capability for safe clinical deployment, requiring the augmentation of current multimodal AI techniques. Our goal in this work is not to add yet another method to the growing toolbox of multimodal AI, but rather to (i) clarify how existing supervised multimodal AI methods can be understood through the lens of uncertainty quantification, and (ii) develop a unified framework that provides valid and interpretable uncertainty estimates for multimodal AI practitioners. We introduce Coracle, a conformalized framework for multimodal AI that seamlessly adapts to early, late, and intermediate fusion. Coracle provides theoretical marginal confidence guarantees and achieves valid finite-sample coverage without relying on distributional assumptions. Through extensive simulations and analyses of diverse biomedical multiview datasets, we demonstrate that Coracle consistently outperforms existing non-conformalized methods in both uncertainty estimation and calibration. By equipping published multiview models with valid, interpretable confidence measures, Coracle advances the development of trustworthy, uncertainty-aware statistical tools for biomedical decision support. Coracle is publicly available as an open-source R package at https://github.com/himelmallick/Coracle.

## 1. Introduction

Multimodal artificial intelligence (AI) has matured over the past decade to the point that foundation models are now routine (Guo et al., 2025; Khan et al., 2025; Li et al., 2024), multimodal endpoints are integral to biomedical research (Acosta et al., 2022; Mallick et al., 2024; Roy et al., 2025; Anceschi et al., 2024; Ding et al., 2022), and the first glimpses of the translational potential of agentic AI have been realized (Swanson et al., 2025; Zou and Topol, 2025; Gao et al., 2024). These advances collectively underscore a paradigm shift from unimodal AI systems that process each data source independently to integrative, multiview AI frameworks that leverage the complementary strengths of diverse modalities (Li and Lock, 2025; Mallick et al., 2025). As a result, researchers are increasingly adopting multiview AI approaches to extract and combine cross-modal information spanning molecular, imaging, clinical, and behavioral domains (Mallick et al., 2017; Tarazona et al., 2021; Xu and McCord, 2022; Argelaguet et al., 2021). This transition has been accelerated by the convergence of large-scale data collection, distributed computing, and probabilistic learning, enabling multimodal AI to move from proof-of-concept studies to real-world applications (Lloyd-Price et al., 2019; Stelzer et al., 2021; Ghaemi et al., 2019; Mallick et al., 2024; Roy et al., 2025; Anceschi et al., 2024; Franzosa et al., 2019).

However, as multimodal models become increasingly central to high-stakes biomedical and clinical decision-making, a crucial question arises: how certain are these models about their predictions (Chakraborti et al., 2025; Kompa et al., 2021)? While supervised multiview integration methods have made impressive strides in accuracy and efficiency, the quantification and communication of uncertainty remain underdeveloped (Roy et al., 2025; Anceschi et al., 2024; Mallick et al., 2024). Many current approaches emphasize point estimates (Franzosa et al., 2019; Stelzer et al., 2021; Ghaemi et al., 2019; Ding et al., 2022), yielding high-performing but often overconfident predictions. Methods that do attempt to quantify predictive uncertainty frequently suffer from poor calibration (Gibson, 2025), leading to uncertainty estimates that fail to achieve reliable frequentist coverage under model misspecification or limited data (Gibson, 2025; Grünwald, 2018; Deliu and Liseo, 2025).

Uncertainty quantification (UQ), when performed rigorously, serves not only as a safeguard for prediction reliability but also as an engine for scientific discovery (Wang et al., 2025; Chakraborti et al., 2025). It highlights cases where data are insufficient, modalities conflict, or additional measurements would be most informative (Bezirganyan et al., 2024; Zhang et al., 2024). Characterizing these uncertainties provides critical insight into which modalities drive confidence, which combinations of views are unstable, and how to prioritize future data collection. As multimodal AI becomes deeply embedded in biomedical research and healthcare, the ability to say “I’m not sure” or “I don’t know” when uncertain is essential for the development of trustworthy and reliable AI systems (López et al., 2025; Kompa et al., 2021).

A number of statistical paradigms attempt to address the need for reliable uncertainty quantification in multimodal settings (Mallick et al., 2025, 2024). Bayesian multimodal approaches provide a natural way to quantify uncertainty by treating model parameters as random variables and propagating posterior distributions through the predictive process (Roy et al., 2025; Anceschi et al., 2024; Mallick et al., 2024). In principle, this framework captures both model variability (epistemic uncertainty) and inherent data noise (aleatoric uncertainty) across multiple modalities. In practice, however, Bayesian multimodal methods often face serious scalability challenges, as posterior inference typically requires computationally intensive procedures or relies on variational approximations that can compromise calibration (Gibson, 2025). Even when posterior inference is computationally feasible, posterior uncertainty may be miscalibrated due to model misspecification or approximate inference, which limits its reliability in complex multimodal applications (Grünwald, 2018). By contrast, frequentist multimodal approaches cannot quantify predictive variability directly and typically require post hoc resampling to produce uncertainty estimates (Ding et al., 2022; Kyung et al., 2010; Stelzer et al., 2021; Franzosa et al., 2019; Ghaemi et al., 2019). Both Bayesian and frequentist paradigms thus lack the finite-sample coverage guarantees necessary to make uncertainty interpretable in multimodal AI systems, and neither extends easily or systematically across early, intermediate, and late fusion paradigms.

In this paper, we introduce Coracle, a conformalized framework for multimodal integration that seamlessly adapts to early, late, and intermediate fusion paradigms (Ding et al., 2022; Mallick et al., 2025). Coracle provides finite-sample marginal coverage guarantees without relying on distributional or parametric assumptions, ensuring statistically valid uncertainty estimates across diverse integration architectures. Through extensive simulation studies and analyses of heterogeneous real-world multiview datasets, we demonstrate that Coracle achieves superior UQ performance metrics relative to existing non-conformalized approaches. The open-source implementation of Coracle is available as an R package at https://github.com/himelmallick/Coracle.

## 2. Coracle - Conformal Prediction for Multiview Learning

In this section, we describe the methodological foundations of our approach. We begin by introducing the multiview learning setup (**Section 2.1**) and reviewing the primary fusion paradigms used in supervised multiview learning (**Section 2.2**). We then introduce the conformal prediction framework (**Section 2.3**) and briefly review split conformal prediction in the single-view setting (**Section 2.4**). Building on these components, we present multiview split conformal prediction, which forms the basis of our proposed Coracle framework (**Section 2.5**), and establish its theoretical guarantees in **Section 2.6**. Finally, we conclude with a brief review of how our method is connected to existing approaches in the literature (**Section 2.7**).

At a high level, Coracle is a model-agnostic framework for uncertainty quantification in multiview prediction and classification (**Fig. 1**). Given any multiview predictive architecture, it partitions the data into training and calibration splits and applies conformal prediction to construct finite-sample valid prediction sets or intervals. By design, this decouples point prediction from uncertainty calibration, allowing existing multiview models to be endowed with rigorous uncertainty guarantees without modifying their internal training procedures. We describe the core methodology underlying Coracle in the sequel.

**Figure 1.**
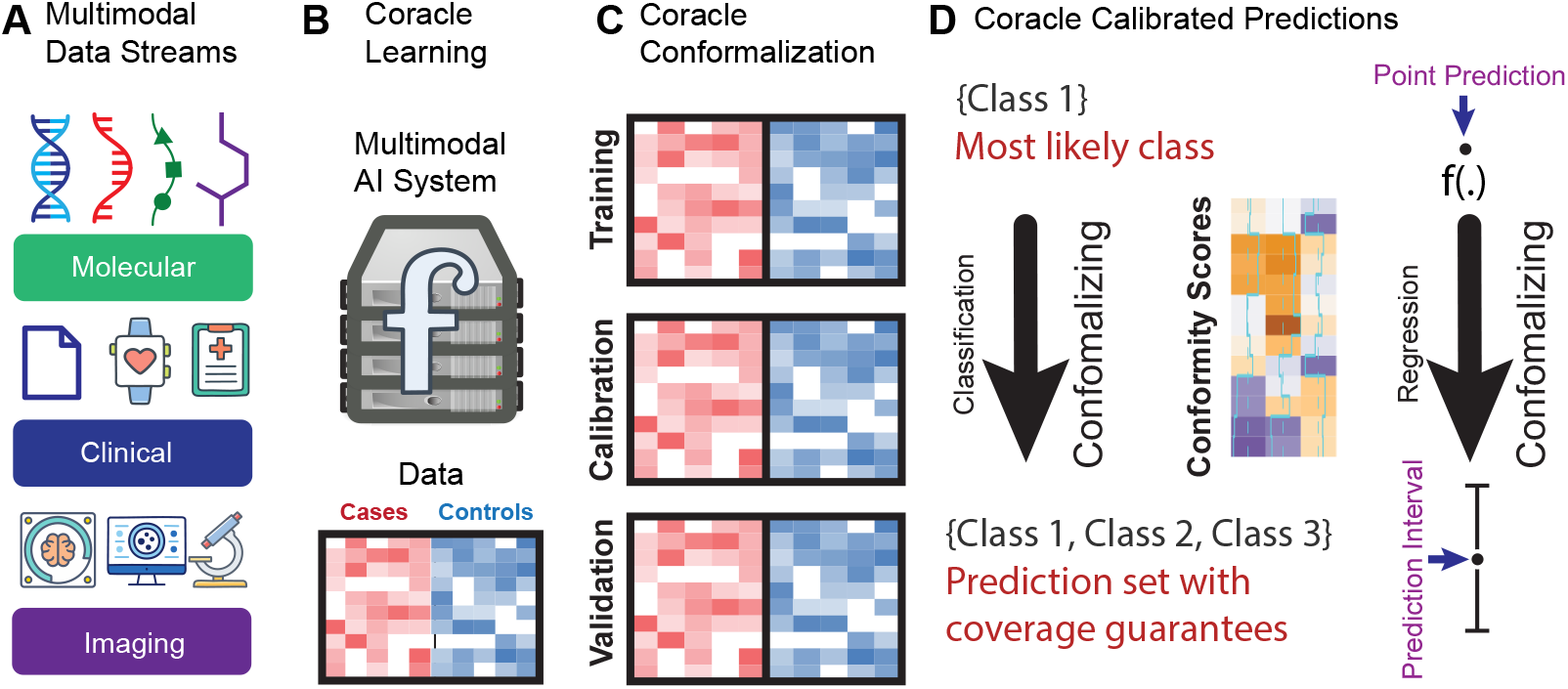
Overview of the Coracle framework for uncertainty-aware multimodal learning. **(A)** Multimodal data streams from molecular, clinical, and imaging sources serve as complementary inputs. **(B)** Coracle learning integrates heterogeneous data views through a unified multiview framework with sequential training, calibration, and validation phases. The predictive function *f* is intentionally left unspecified, allowing it to represent any multimodal learning architecture (e.g., early, intermediate, or late fusion; deep or shallow models), which makes the framework model-agnostic and broadly applicable. **(C)** Coracle conformalization applies conformal prediction using a held-out calibration set to compute conformity scores that quantify predictive uncertainty. **(D)** Coracle calibrated predictions produce uncertainty-aware outputs, including classification prediction sets and regression prediction intervals with finite-sample coverage guarantees.

### 2.1 Multiview Learning Setup

We consider a supervised learning setting where each sample consists of a response *Y*_*i*_ ∈ 𝒴 and *K* distinct input views 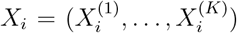, with 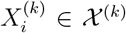 representing the *k*-th view. The goal is to learn a predictive function

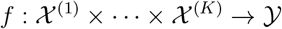

that maps multiview inputs to the output space. The training dataset is denoted as 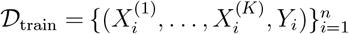, where all samples are assumed to be i.i.d. from an unknown distribution over 𝒳 ^(1)^ × · · · × 𝒳 ^(*K*)^ × 𝒴. The structure of *f* depends on the fusion paradigm employed, which governs how multiview inputs are processed and integrated. For conformal prediction, we further assume that the calibration and test samples are exchangeable with the training data.

### 2.2 Multiview Fusion Paradigms

Multiview learning addresses the problem of integrating multiple heterogeneous data sources (or views) that provide complementary information about a common underlying phenomenon. Depending on when and how integration is performed, multiview methods are typically categorized into three paradigms: *early fusion, late fusion*, and *intermediate fusion* (Li et al., 2016). These paradigms differ in representational flexibility, interpretability, computational cost, and their ability to capture cross-view interactions (Mallick et al., 2025). The choice of fusion strategy determines how the input views are combined and, consequently, how the predictive function *f* is constructed.

#### 2.2.1 Early Fusion

In early fusion, the views are concatenated to form a single composite feature vector,

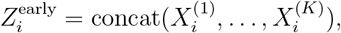

and a single predictor is trained on this fused representation. The resulting prediction function is *f*_early_ : *Ƶ*^early^ → 𝒴, where *Ƶ*^early^ = 𝒳 ^(1)^ × · · · × 𝒳 ^(*K*)^. This approach assumes that cross-view interactions can be adequately captured at the feature level. Standard learning algorithms (e.g., random forests (Breiman, 2001), Bayesian additive regression trees (BART) (Chipman et al., 2010), or neural networks (Goodfellow et al., 2016; Papadopoulos et al., 2007)) may be applied directly to the fused representation, possibly after view-specific dimensionality reduction such as PCA or autoencoders (Vincent et al., 2010). While early fusion is conceptually simple and compatible with off-the-shelf learning pipelines, it may obscure view-specific structure and can become computationally challenging when the views are high-dimensional or poorly aligned (Anceschi et al., 2024; Roy et al., 2025).

#### 2.2.2 Late Fusion

In late fusion, each view is modeled independently, and predictions are combined only at the decision level. Given views ***X***^(1)^, …, ***X***^(*K*)^, separate predictors *f* ^(*k*)^ : 𝒳 ^(*k*)^ → 𝒴 are trained, yielding predictions *f* ^(*k*)^(***X***^(*k*)^). A fusion function *ϕ* : 𝒴^*K*^ → 𝒴 then aggregates these outputs to form the final predictor,

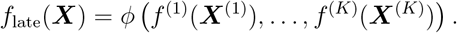

Common aggregation rules include averaging, voting, stacking, and weighted ensembles (Van der Laan et al., 2007). Late fusion offers modularity and interpretability, as each view-specific model can be inspected independently. However, because the views are modeled in isolation, this paradigm may fail to capture synergistic cross-view interactions, and performance can be sensitive to the choice of aggregation rule (Anceschi et al., 2024; Roy et al., 2025).

#### 2.2.3 Intermediate Fusion

Intermediate fusion lies between early and late fusion by allowing joint modeling of multiple views while preserving view-specific structure. A representative example is the cooperative learning framework (Ding et al., 2022), which encourages agreement across view-specific predictors through regularization and minimizes the objective

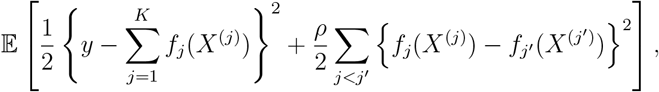

where *ρ* ⩾ 0 controls the strength of the agreement penalty. When *ρ* = 0, the views are effectively modeled independently, while as *ρ* → ∞, the predictors are forced to agree, recovering a behavior closer to late fusion. More generally, intermediate fusion constructs a joint predictor

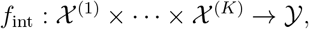

learned using architectures that explicitly retain view structure. This paradigm enables cross-view interaction while avoiding both full feature-level concatenation and complete view-wise decoupling (Anceschi et al., 2024; Roy et al., 2025). Across all three paradigms, we denote the resulting predictor generically by *f*, with the understanding that its internal construction depends on the chosen fusion strategy, while its interface remains *f* (*X*^(1)^, …, *X*^(*K*)^) ∈ 𝒴.

### 2.3 Conformal Prediction

*A General Framework for Uncertainty Quantification* Conformal prediction (CP) provides a general framework for constructing prediction sets or intervals with finite-sample coverage guarantees (Angelopoulos and Bates, 2021; Shafer and Vovk, 2008). Given a trained predictor *f*, CP quantifies uncertainty by calibrating conformity scores on a held-out calibration set under the assumption of exchangeability between calibration and test points (Angelopoulos and Bates, 2021; Shafer and Vovk, 2008). Let *S*_*i*_ = score(*Y*_*i*_, *f* (*X*_*i*_)) denote a conformity score measuring the discrepancy between the prediction and the observed response. Given calibration scores 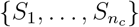, the (1−*α*)-level conformal prediction set for a test input *X*_*n*+1_ is

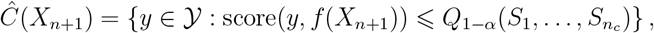

where *Q*_1−*α*_ denotes the (1−*α*)(1+1*/n*_*c*_)-quantile of the calibration scores. This construction applies uniformly across fusion paradigms by substituting the appropriate predictor *f*, i.e., *f*_early_, *f*_late_, or *f*_int_. To formalize the Coracle framework, let 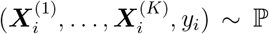, for *i* = 1, …, *n*, be i.i.d. observations from a distribution ℙ on 𝒳 ^(1)^ × · · · × 𝒳 ^(*K*)^ × 𝒴. The objective is to construct a prediction set or interval *Ĉ*_*n*_(·) such that, for a new input,

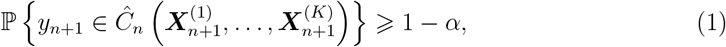

for a prescribed level 1 − *α*.

### 2.4 Single-view Split Conformal Prediction

We briefly review split conformal prediction (SCP) in the single-view setting, which serves as a building block for our multiview construction. SCP is a model-agnostic procedure for constructing prediction intervals with finite-sample marginal coverage under the assumption of exchangeability (Shafer and Vovk, 2008; Angelopoulos and Bates, 2021). Suppose we observe i.i.d. data 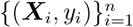, where ***X***_*i*_ ∈ 𝒳 ⊂ ℝ^*p*^ and *y*_*i*_ ∈ ℝ. The objective is to construct a prediction interval *Ĉ*_*n*_(***X***_*n*+1_) such that

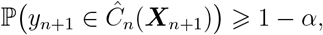

for a given *α* ∈ (0, 1). SCP proceeds as follows. The data are split into a training set of size *n*_1_ and a calibration set of size *n*_2_ = *n* − *n*_1_. A predictor 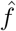 is fit on the training set, and conformity scores (here, absolute residuals) are computed on the calibration set,

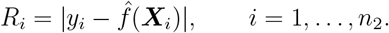

Let 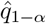 denote the ⌈(1 − *α*)(*n*_2_ + 1)⌉-th order statistic of 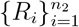. The resulting prediction interval is

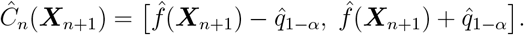

This construction satisfies the marginal coverage guarantee in Equation (1) for *d* = 1 under the sole assumption of exchangeability. While we focus on regression here, the same framework extends to classification and other prediction tasks through appropriate choices of conformity scores (Shafer and Vovk, 2008; Angelopoulos and Bates, 2021), as detailed in **Algorithm 1**.

### 2.5 Multiview Split Conformal Prediction

Having described single-view SCP, we now show how it extends naturally to multiview learning. We assume that each observation consists of a response *Y* ∈ 𝒴 and *K* input views *X* = (*X*^(1)^, …, *X*^(*K*)^), where *X*^(*k*)^ ∈ 𝒳 ^(*k*)^. A predictor *f* maps from the product space 𝒳 ^(1)^ × · · · × 𝒳 ^(*K*)^ to 𝒴. The conformalization step is identical across fusion paradigms and proceeds exactly as in single-view SCP: given a fused predictor *f*, SCP is applied to its residuals. The paradigms differ only in the construction of *f*. We consider the three standard fusion paradigms below.

#### 2.5.1 Early Fusion

In early fusion, the views are first combined at the feature level, and a single predictor is trained on the resulting representation. Specifically, we form the concatenated feature vector

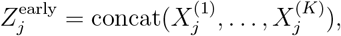

and train a predictor *f*_early_ on *Z*^early^. Conformity scores are then computed in the usual way,

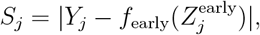

and SCP is applied to obtain the prediction set *Ĉ*(*X*_*n*+1_).

#### 2.5.2 Late Fusion

In late fusion, each view is modeled separately, and their predictions are combined at the decision level. Concretely, we train view-specific predictors *f* ^(1)^, …, *f* ^(*K*)^ and aggregate their outputs through a fusion function *ϕ*,

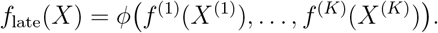

In principle, conformalization can be applied either before or after fusion. One may conformalize each *f* ^(*k*)^ separately and then aggregate the resulting sets (modality-level CP), or conformalize the fused predictor directly (fusion-level CP). We adopt the latter strategy, computing residuals

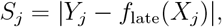

and applying SCP to obtain a single, unified prediction set.

#### 2.5.3 Intermediate Fusion

Intermediate fusion lies between these two extremes: the views are processed jointly, but their structure is preserved within the model. This includes, for example, cooperative learning architectures (Ding et al., 2022). In this case, we train a joint predictor *f*_int_ on (*X*^(1)^, …, *X*^(*K*)^) and compute conformity scores as

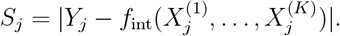

Once again, SCP is applied without modification.

In all three cases, Coracle treats the multiview predictor as a black box and applies the same conformalization step to its residuals (**Algorithm 2**). This viewpoint makes clear that the proposed framework is not tied to any specific fusion architecture, but instead provides a uniform mechanism for equipping a wide range of multiview models with finite-sample valid prediction intervals.

### 2.6 Theoretical Guarantees

CP guarantees marginal coverage at the nominal level 1 − *α*, provided that the calibration and test points are exchangeable and that the function *f* used to compute conformity scores is fixed and trained independently of the calibration set (Angelopoulos and Bates, 2021; Shafer and Vovk, 2008). These conditions are satisfied for all fusion paradigms described above. In particular, for any fusion paradigm and any such function *f*,

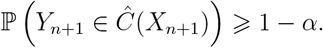

That is, regardless of the underlying fusion architecture, the prediction set *Ĉ*(*X*_*n*+1_) constructed by Coracle contains the true response with probability at least 1 − *α* for a new test point, without any distributional assumptions beyond exchangeability. The Coracle framework, therefore, extends conformal prediction to multiview learning while preserving finite-sample validity across a broad class of models and fusion paradigms (**Fig. 1**).

### 2.7 Relationship with Existing Methods

In this section, we situate our approach within the existing literature on multiview conformal methods. While single-view conformal prediction has seen broad methodological development and diverse applications across many areas of machine learning and statistics (Xu et al., 2024; Sun et al., 2024; Angelopoulos and Bates, 2021; Shafer and Vovk, 2008), its use in multiview or multimodal learning remains extremely limited (Serra et al., 2019). Currently, only a limited number of methods have been proposed for conformal prediction in multiview or multimodal settings (Garcia-Ceja, 2024; Rivera et al., 2024; Bose et al., 2024). Most published methods are confined to classification problems and focus on constructing prediction sets for discrete labels, with limited applicability to continuous outcomes or to disparate fusion architectures. In contrast, continuous outcomes enable a more informative characterization of uncertainty through interval width, empirical coverage, and calibration, since the inferential target is real-valued and prediction sets naturally take the form of intervals. Building on split conformal prediction, our framework delivers finite-sample valid prediction intervals for early-, intermediate-, and late-fusion models across a broad class of outcomes (**Fig. 1**), thereby unifying and substantially extending the current sparse landscape of multiview conformal methods.

The Coracle framework is implemented as an open-source R package and is publicly available at https://github.com/himelmallick/Coracle. The package provides a unified interface for conformalizing early-, intermediate-, and late-fusion multiview predictors, together with tools for data splitting, calibration, and construction of prediction intervals. The implementation is designed to integrate with standard machine learning workflows. We anticipate that this software will serve as a general-purpose resource for the community, facilitating the development, benchmarking, and deployment of uncertainty-aware multiview learning methods.

## 3. Simulation Studies

### 3.1 Simulation Design

We consider two complementary classes of simulation settings to evaluate conformal prediction in multiview learning: (i) *data-driven simulations* based on realistic multi-omics covariance structures derived from The Cancer Genome Atlas (TCGA; (Weinstein et al., 2013)) data using *InterSIM* (Chalise et al., 2016), and (ii) *factor model-based simulations* that explicitly introduce latent shared structure across data views.

#### 3.1.1 Common Setup

For the data-driven simulations, we use *InterSIM* (Chalise et al., 2016) to generate three interconnected data modalities: gene expression, DNA methylation, and protein abundance, based on ovarian cancer data from TCGA. Each simulated dataset contains 658 features spanning three omics layers (131 gene expression, 367 methylation, and 160 protein features). We generate 25 replicate datasets and split each into 70% training and 30% test sets. We consider two sample sizes, *n* = 200 and *n* = 500, and vary sparsity levels across 10% and 50% active features, representing sparse and dense regimes. Outcomes are generated according to

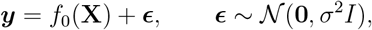

where *σ*^2^ is calibrated to achieve a desired signal-to-noise ratio (SNR), defined as 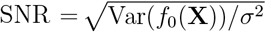. We consider SNR = 10 (high), SNR = 5, 3 (moderate), and SNR = 1 (low).

#### 3.1.2 Data-Driven Simulations: Mixed Linear and Nonlinear Effects

Within the *Inter-SIM* framework, we generate outcomes using

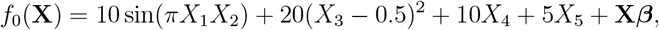

where (*X*_1_, …, *X*_5_) are randomly selected features and the first five coefficients of ***β*** are set to zero (Mallick et al., 2024). Only a fixed proportion (10% or 50%) of the remaining coefficients are nonzero, drawn from a log-normal distribution calibrated using the Splatter framework (Zappia et al., 2017; Mallick et al., 2022) to yield realistic log_2_ fold-changes in the range (−3, 3). This setting combines structured nonlinear signal components with highdimensional sparse linear effects under a realistic multi-omics correlation structure (Mallick et al., 2022, 2024).

### 3.2 Factor model-Based Simulations: Latent Shared Structure

To assess performance under explicit shared latent structure across views, we additionally consider a multiview factor model (Anceschi et al., 2024; Ding et al., 2022; Samorodnitsky et al., 2024). Let 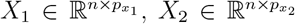, and 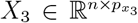 denote three data views. We first generate all features independently from MVN(0, *I*_*n*_). Next, we generate *p*_*u*_ latent factors 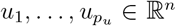 independently from MVN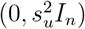 and inject shared structure into each view via

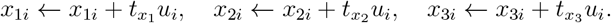

Letting 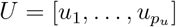, the outcome is generated as

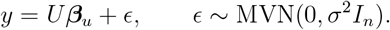

This setting creates a scenario in which the predictive signal is driven primarily by latent factors shared across views rather than directly by observed features.

#### 3.2.1 Choice of Baseline Methods and Evaluation Criteria

While many supervised multi-view learning methods exist, we deliberately focus on *IntegratedLearner* (Mallick et al., 2024) for early and late fusion and *BayesCOOP* (Roy et al., 2025) for intermediate fusion. These methods represent two of the very few Bayesian multiview learning frameworks that natively provide uncertainty quantification through posterior inference, and *IntegratedLearner* is widely used as a principled and interpretable baseline for multiview integration (Mansoor et al., 2024; Lac et al., 2024; Sherwani et al., 2025; Hernández-Lemus and Ochoa, 2024; Li and Lock, 2025; Zhang et al., 2025; Mallick et al., 2025).

Briefly, *IntegratedLearner* is a Bayesian ensemble framework for multi-omics prediction and classification that fits view-specific machine learning models (with BART as the default, while also supporting learners such as random forests and XGBoost) and combines them through a meta-learner to improve overall predictive performance and interpretability (Mallick et al., 2024). *BayesCOOP* is a Bayesian group spike-and-slab approach that integrates multiview predictions within a Bayesian bootstrap framework, encouraging agreement across views while quantifying uncertainty (Roy et al., 2025). Together, they span the three principal fusion paradigms (early, intermediate, and late), enabling a controlled and principled comparison of conformalization across fusion strategies while keeping the underlying learning frameworks fixed. Across all experiments, we report empirical coverage probabilities at multiple quantiles to assess validity, and for the real-data analyses, we additionally report the average width of the prediction intervals to assess practical efficiency (**Section 4**).

#### 3.2.2 Simulation Results

Several observations are in order. Across all simulation settings considered, Coracle delivers robust uncertainty quantification, consistently achieving near-nominal coverage for early, intermediate, and late fusion models. In contrast, the corresponding non-conformal baselines (*IntegratedLearner* and *BayesCOOP*) exhibit noticeable under-coverage, particularly in low-signal and small-sample regimes (**Figs. 2–3**). These results underscore the necessity of the conformal calibration step when the signal is weak or the available training data are limited. Additional simulations exploring moderate signal-to-noise ratios and alternative late-fusion base learners, including random forests (Breiman, 2001; Liaw and Wiener, 2002) and XGBoost (Chen and Guestrin, 2016) (**Figs. S1–S7**), further confirm the robustness of Coracle across a broad range of settings.

**Figure 2.**
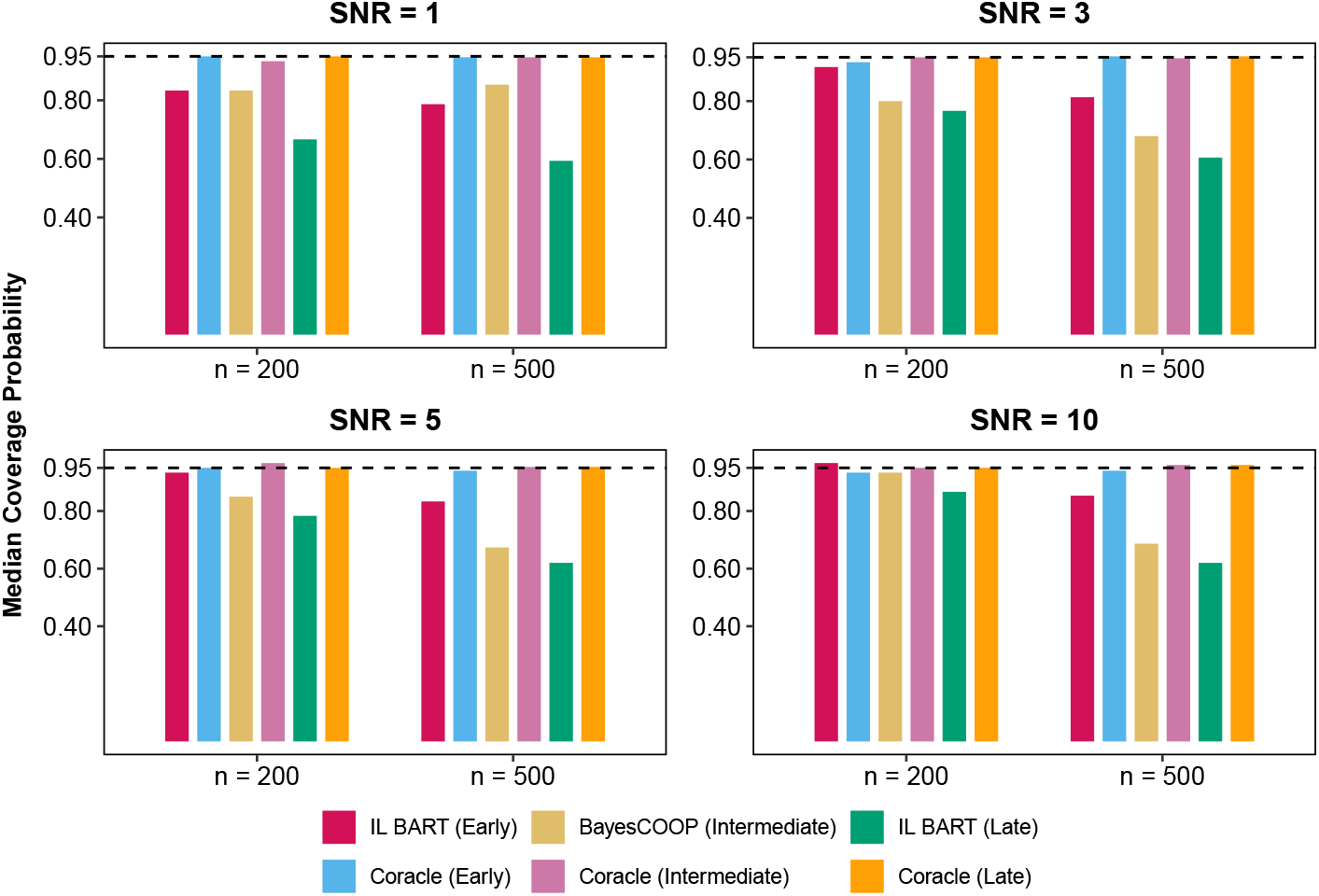
Coracle significantly outperforms published methods in predictive UQ in factor model–based simulations (Anceschi et al., 2024; Ding et al., 2022). Coverage performance for Coracle and published non-conformal baselines is evaluated under four signal-to-noise ratios (SNR = 1, 3, 5, 10) and two sample sizes (*n* = 200, 500). Here, *IL BART* denotes *IntegratedLearner* (Mallick et al., 2024) with BART (Chipman et al., 2010) as the base learner. *BayesCOOP* (Roy et al., 2025) represents the intermediate fusion baseline. For each configuration, median coverage probabilities are computed using completely held-out test data and summarized over 25 replications. Coracle maintains high and stable coverage across all scenarios and across early, intermediate, and late fusion architectures, demonstrating robust uncertainty quantification, whereas alternative non-conformalized fusion methods show notable deviations from nominal coverage, especially in low-signal and low-sample regimes.

**Figure 3.**
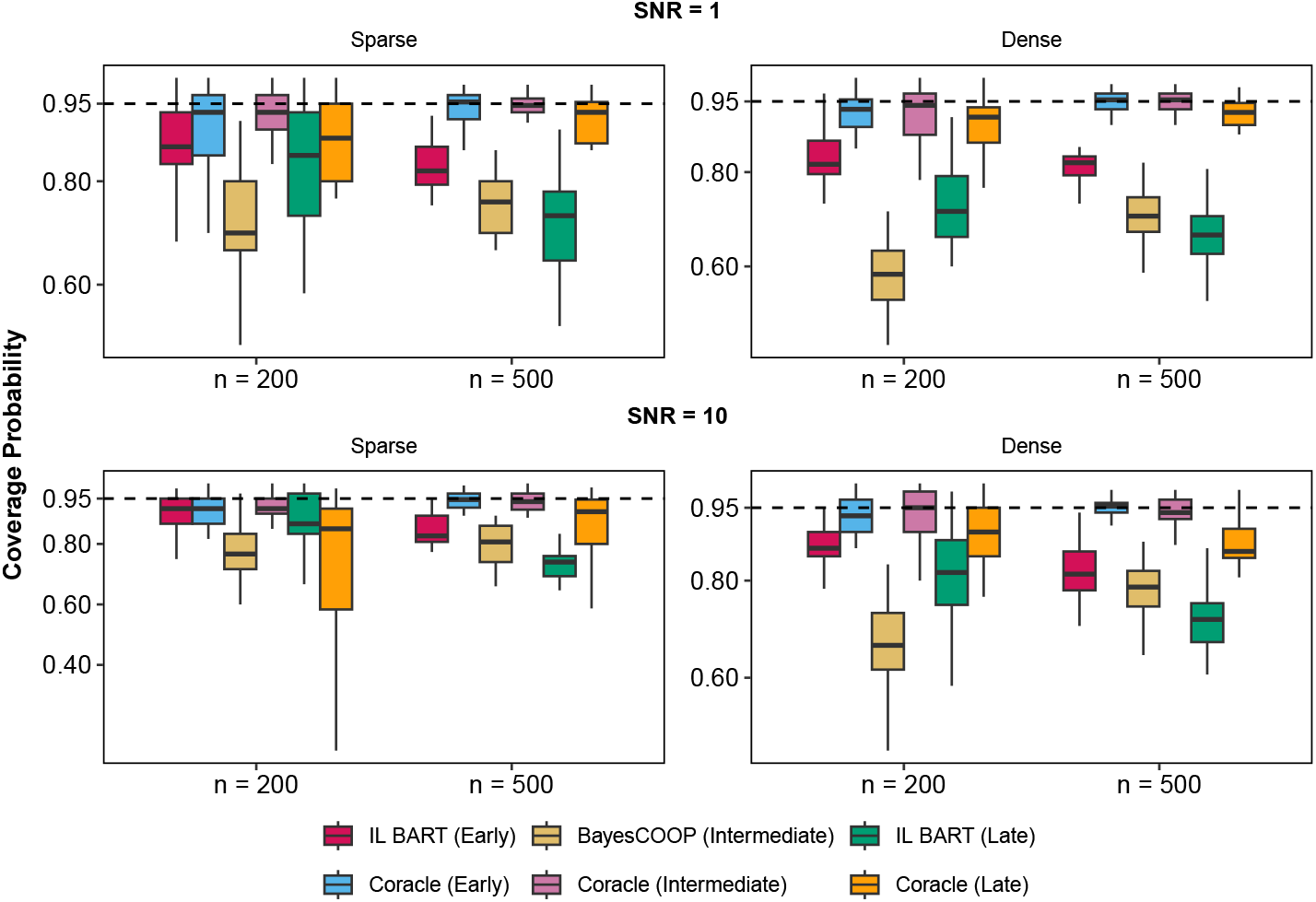
Coracle significantly outperforms published methods in predictive UQ in data-driven, non–factor model–based simulations with highly nonlinear effect sizes (Mallick et al., 2024; Chalise et al., 2016). Coverage performance for Coracle and published non-conformal baselines is evaluated under two signal-to-noise ratios (SNR = 1 and SNR = 10), two sample sizes, and both sparse and dense coefficient regimes. Here, *IL BART* denotes *IntegratedLearner* (Mallick et al., 2024) with BART (Chipman et al., 2010) as the base learner. *BayesCOOP* (Roy et al., 2025) represents the intermediate fusion baseline. For each configuration, median coverage probabilities are computed using completely held-out test data and summarized over 25 replications. Coracle maintains high and stable coverage across all scenarios and across early, intermediate, and late fusion architectures, demonstrating robust UQ, whereas non-conformal fusion counterparts show substantial under-coverage, particularly in low-signal and small-sample configurations.

## 4. Real Data Analyses

### 4.1 Pregnancy Multimodal Datasets

We consider two longitudinal multi-omics datasets, StelzerDOS and StelzerEGA, originating from a recent study conducted at Stanford University (Stelzer et al., 2021). To investigate the temporal trajectory of pregnancy-related immunologic changes, 53 pregnant women receiving routine antepartum care at Lucile Packard Children’s Hospital (Stanford, CA, USA) were enrolled, and venous peripheral blood samples were collected during the final 100 days of pregnancy. This yielded 2,757 features per sample across two molecular layers, namely CyTOF and proteomics, for both datasets.

The StelzerDOS dataset uses time to spontaneous labor as the outcome of interest, defined as the day of admission for spontaneous labor, characterized by contractions occurring at least every five minutes, lasting more than one minute, and associated with cervical changes. The StelzerEGA dataset, in contrast, focuses on gestational age as the outcome variable, determined according to the best obstetrical estimate as recommended by the American College of Obstetricians and Gynecologists (Ghaemi et al., 2019). Both datasets include a held-out test set consisting of 8 subjects. For simplicity of analysis, we consider only the baseline observations.

### 4.2 Inflammatory Bowel Disease Multimodal Dataset

We also analyze a dataset from the Prospective Registry for IBD Study at Massachusetts General Hospital (PRISM) (Franzosa et al., 2019; Mallick et al., 2019), which contains 9,171 quality-controlled features derived from two molecular layers and serves as a benchmark for evaluating the performance of Coracle. The PRISM cohort is a cross-sectional study designed to characterize the gut metabolic landscape and microbiome composition in individuals with IBD and includes 155 participants: 68 with Crohn’s disease (CD), 53 with ulcerative colitis (UC), and 34 non-IBD controls. Stool samples from all participants underwent metagenomic sequencing to quantify microbial taxonomic composition and functional potential. In parallel, multi-omics profiling was performed using four liquid chromatography–tandem mass spectrometry (LC–MS) platforms measuring polar metabolites, lipids, free fatty acids, and bile acids.

For our analysis, we construct the dysbiosis score following Lloyd-Price *et al*. (Lloyd-Price et al., 2019). Briefly, the dysbiosis score is defined as the median Bray–Curtis dissimilarity between a given test sample and a healthy reference population (non-IBD), and serves as the outcome variable of interest. In addition, the dataset includes an independent validation set of 65 individuals, enabling assessment of out-of-sample predictive performance.

### 4.3 Summary of Findings in Real Data

Across the three real-world multiview datasets, we first assess the predictive accuracy of the underlying base learners before conformalization (**Fig. 4**). The mean squared prediction error (MSPE) results show that early and intermediate fusion models generally achieve the lowest errors, while late fusion performs slightly worse. This confirms that the base predictors targeted for conformalization are already competitive, and that subsequent gains arise primarily from uncertainty calibration rather than from changes in point prediction accuracy.

**Figure 4.**
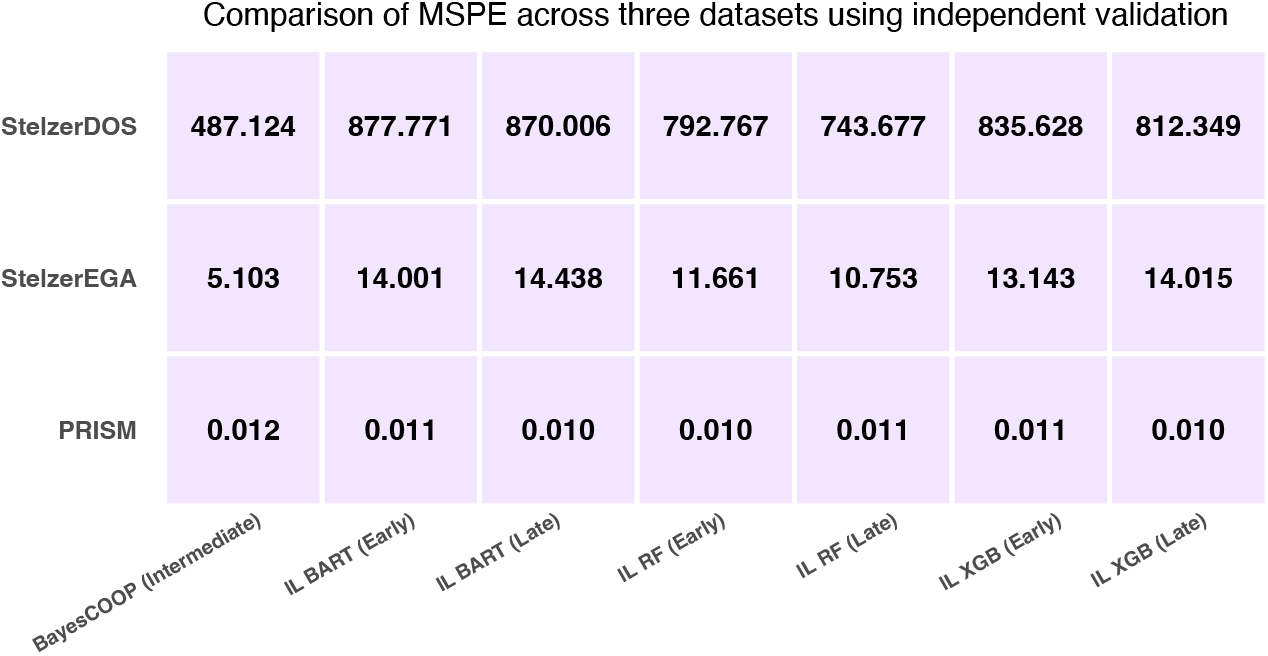
Mean squared prediction error (MSPE) comparison across three datasets for various fusion paradigms. MSPE is evaluated using independent validation data for three studies (PRISM (Franzosa et al., 2019), StelzerEGA (Stelzer et al., 2021), and StelzerDOS (Stelzer et al., 2021)). Representative methods from each fusion paradigm (early, intermediate, and late) are shown, illustrating that the underlying base models targeted for conformalization provide competitive baseline predictions prior to UQ calibration. Here, *IL* denotes *IntegratedLearner* (Mallick et al., 2024) with different base learners (Van der Laan et al., 2007): *IL BART* uses BART (Chipman et al., 2010), *IL RF* uses random forest (Breiman, 2001; Liaw and Wiener, 2002), and *IL XGB* uses XGBoost (Chen and Guestrin, 2016) (each shown under early and late fusion). *BayesCOOP* (Roy et al., 2025) represents the intermediate fusion baseline.

After assessing predictive performance on the held-out test subjects, we compare subject-level prediction intervals across fusion paradigms. In the pregnancy datasets, we observe that Coracle-conformalized models achieve near-nominal coverage across all subjects (**Figs. 5; S8**), while non-conformal baselines sometimes fail to contain the true outcome. Early and intermediate fusion models produce comparatively narrow intervals, reflecting efficient integration of proteomics and CyTOF features, whereas late fusion yields wider intervals and less reliable coverage. A similar pattern is observed in the PRISM IBD dataset (**Fig. 6**). Early and intermediate fusion models benefit substantially from conformalization, showing clear improvements toward nominal coverage. In contrast, late fusion models perform comparatively poorly at baseline, and while Coracle still offers improvement, the gains are more limited in this dataset, hinting at poor calibration despite good baseline predictive performance.

**Figure 5.**
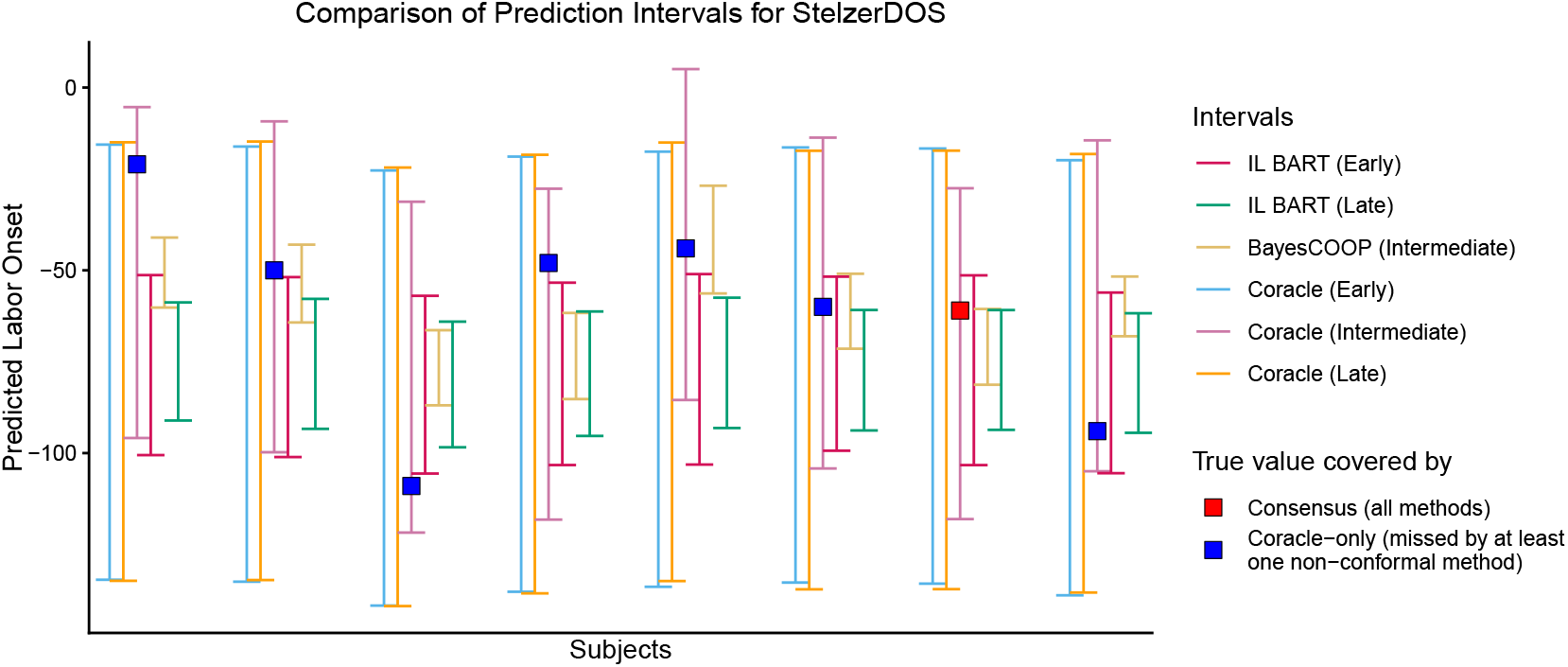
Prediction interval comparison for the StelzerDOS dataset(Stelzer et al., 2021). Subject-level prediction intervals for labor onset time are shown for representative early, intermediate, and late fusion base models and their corresponding Coracleconformalized versions, evaluated on completely independent validation data. Here, *IL BART* denotes *IntegratedLearner* (Mallick et al., 2024) with BART (Chipman et al., 2010) as the base learner (shown under early and late fusion), and *BayesCOOP* (Roy et al., 2025) denotes the Bayesian cooperative learning model used as the intermediate fusion baseline. For each subject, the true value is indicated, along with markers distinguishing cases in which all methods cover the true value from those in which Coracle achieves valid coverage but at least one non-conformal approach fails to do so.

**Figure 6.**
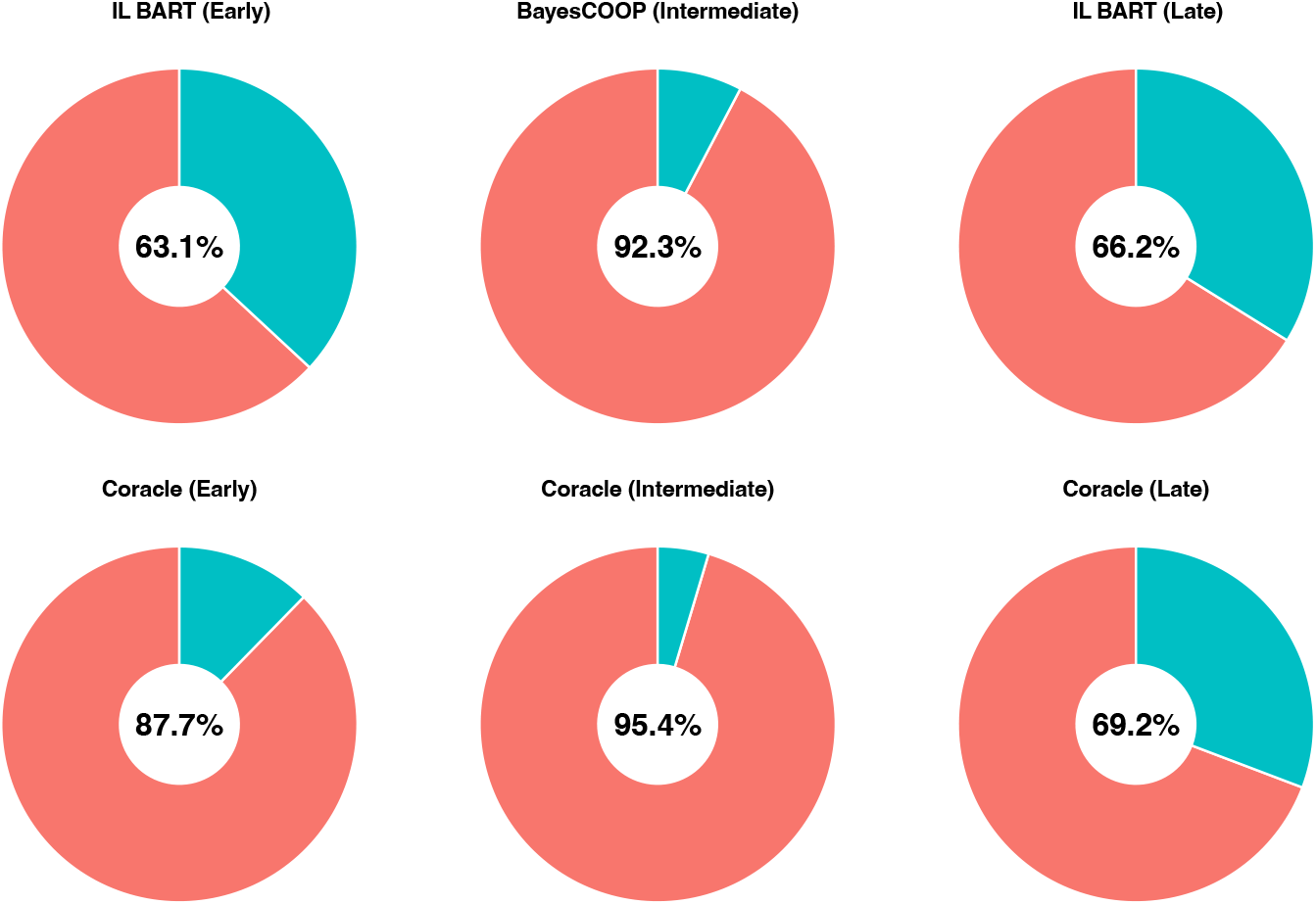
Coverage comparison for the PRISM dataset (Franzosa et al., 2019) across fusion paradigms. Coverage is evaluated on completely independent validation data for early, intermediate, and late fusion base models and their corresponding Coracle-conformalized versions. Here, *IL BART* denotes *IntegratedLearner* (Mallick et al., 2024) with BART (Chipman et al., 2010) as the base learner (used for early and late fusion), and *BayesCOOP* (Roy et al., 2025) denotes the Bayesian cooperative learning model used as the intermediate fusion baseline. Results for the base (non-conformal) models are shown in the top row, and the corresponding Coracle-conformalized results are shown in the bottom row. Early and intermediate fusion models benefit substantially from conformalization, showing clear improvements toward nominal coverage. In contrast, late fusion models perform comparatively poorly at baseline, and while Coracle still offers improvement, the gains are more limited in this dataset, hinting at poor calibration despite good baseline predictive performance.

## 5. Discussion

In this work, we introduce Coracle, a conformal framework for multimodal AI that provides valid, finite-sample UQ across early, intermediate, and late fusion paradigms. Through extensive simulation studies and real-data analyses, we demonstrated that Coracle not only improves predictive calibration but also enhances interpretability by enabling multiview models to communicate when predictions are uncertain. This capability is particularly valuable in biomedical contexts, where decisions must balance predictive accuracy with accountability and patient safety. In such settings, decisions based on a single, overconfident estimate derived from noisy multimodal data can harm patients, underscoring the need for uncertainty quantification and calibration in multimodal AI (Kompa et al., 2021).

Methodologically, Coracle’s conformal design provides a principled yet scalable approach to uncertainty in multimodal learning. It avoids the computational and calibration challenges of Bayesian methods and the ad hoc adjustments of frequentist approaches, while still delivering formal coverage guarantees. This unifies flexibility, scalability, and statistical validity in a single, model-agnostic framework. Beyond these methodological advantages, Coracle contributes to a broader re-framing of multimodal AI as a system that must be aware of its own limitations. Traditional multimodal integration pipelines typically optimize for accuracy, often overlooking whether predictions are stable, reproducible, or trustworthy. By embedding UQ directly into the integration process, Coracle enables practitioners to identify data regimes where predictions are unstable, modalities that contribute dispropor-tionately to uncertainty, and cases where collecting additional data could meaningfully reduce prediction error. This uncertainty-aware perspective shifts the goal of multimodal learning from producing a single best prediction to generating trustworthy predictions, accompanied by transparent confidence measures that can guide downstream inference.

Several avenues remain for future work. First, Coracle currently guarantees marginal coverage across samples but does not ensure group-conditional or modality-conditional coverage, meaning that certain clinically relevant subpopulations or data regimes may still be under-covered. Developing stratified or locally adaptive conformalization strategies could improve fairness and reliability in heterogeneous biomedical settings. Second, while Coracle assumes exchangeability between calibration and test data, real-world multimodal studies often involve distribution shift across cohorts, platforms, or clinical sites. More explicitly integrating epistemic uncertainty or shift-aware diagnostics into the conformal calibration process could improve robustness and enhance the detection of out-of-distribution or low-confidence cases (Liu et al., 2024; Gibbs and Candes, 2021; Tibshirani et al., 2019). Finally, extending Coracle to temporal, spatial, hierarchical, and transfer learning settings (Xu and McCord, 2022; Tian and Feng, 2023; Suder et al., 2025) would further broaden its applicability to longitudinal, cross-study, and high-resolution biomedical analyses.

In summary, Coracle advances the current state of multimodal AI by combining conformal prediction with flexible multimodal machine learning architectures, enabling robust, interpretable, and distribution-free uncertainty quantification across diverse data fusion paradigms. As multimodal AI becomes increasingly central to biomedical and translational research, such frameworks will be crucial to ensure that predictive models are not only accurate but also self-aware -capable of expressing calibrated uncertainty that supports a responsible, evidence-based clinical decision process.

## Code and Data Availability

An open-source implementation of the Coracle framework is available as an R package at https://github.com/himelmallick/Coracle. The public multi-omics datasets used in this study are publicly available and can be obtained from the `IntegratedLearner` repository (Mallick et al., 2024) at https://github.com/himelmallick/IntegratedLearner.

## Algorithm 1

Coracle for single-view split conformal prediction

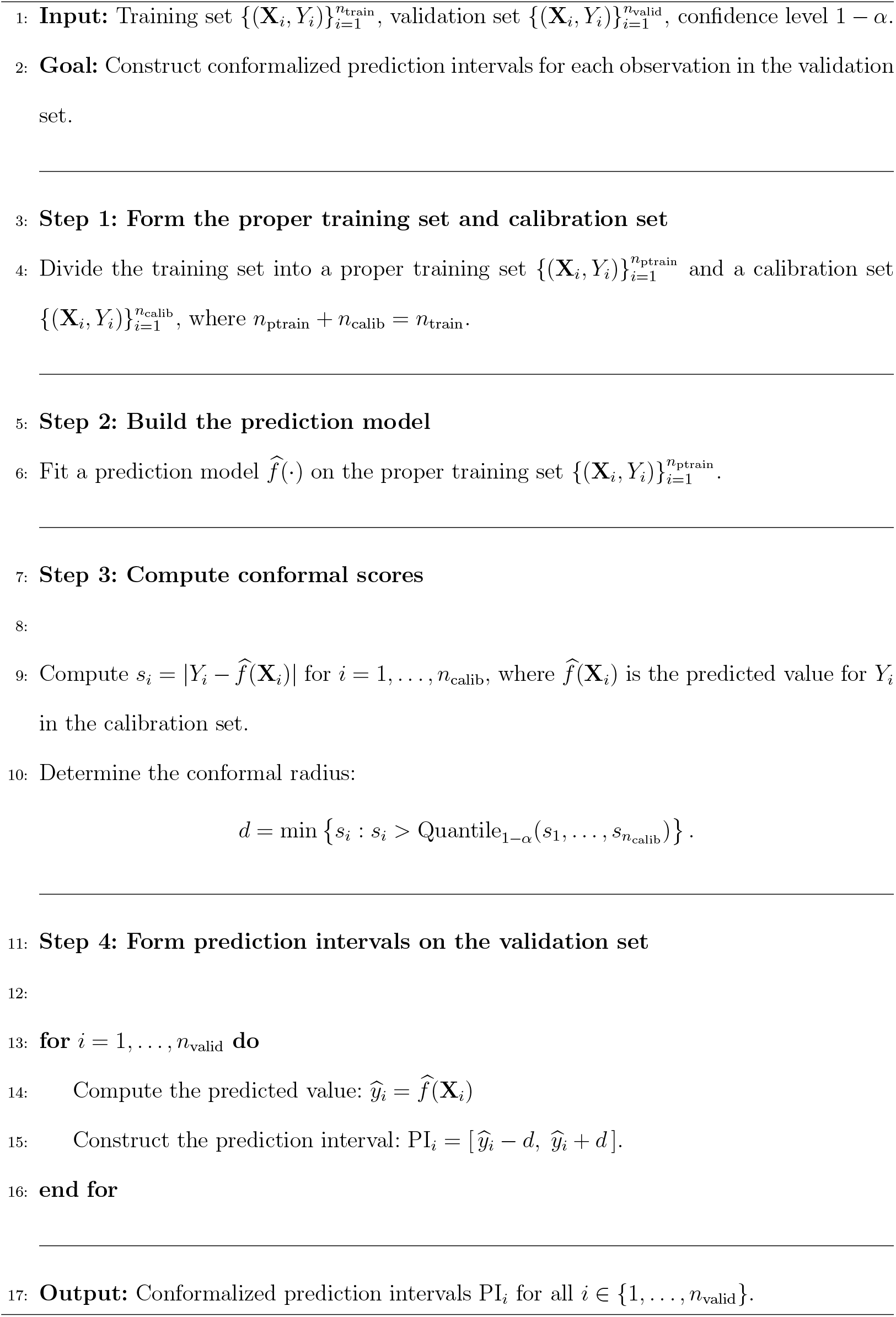

## Algorithm 2

Coracle for multi-view split conformal prediction

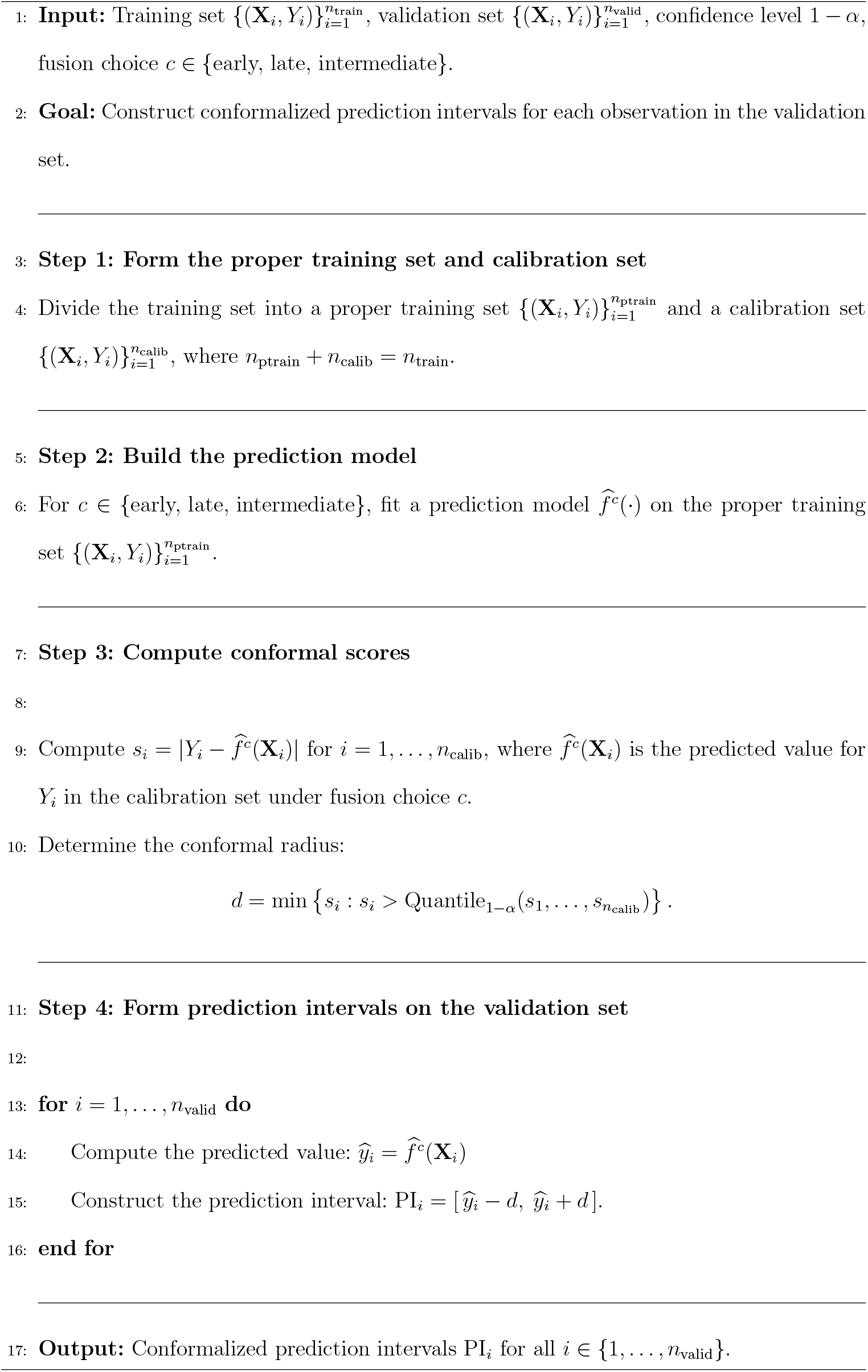

