## Supplementary Materials for "Conformalized Multiview Learning"

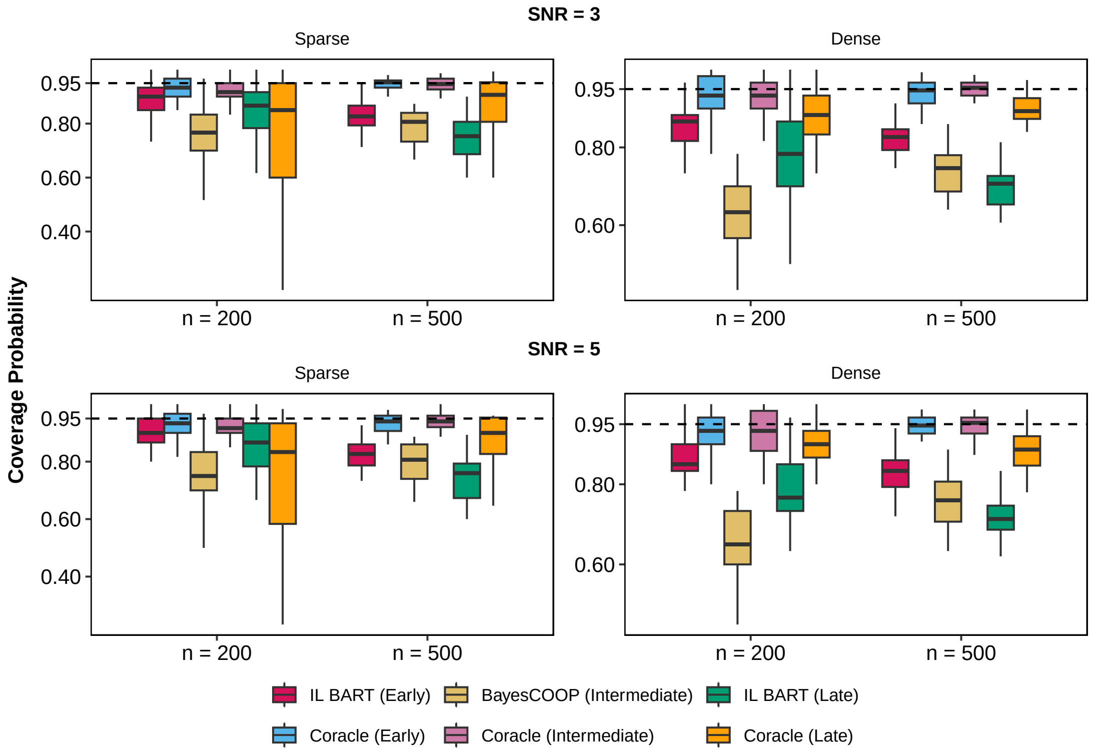


Figure S1: **Data-driven, non-factor model–based simulations with BART as base learner for late fusion in moderate SNR settings**. Coverage performance for Coracle and published non-conformal baselines is evaluated under two signal-to-noise ratios (SNR $= 3$ and $5$), two sample sizes ($n = 200, 500$), and both sparse and dense coefficient regimes. Here, *IL BART* denotes *IntegratedLearner* (Mallick et al., 2024) with BART as the base learner (Chipman et al., 2010), and *BayesCOOP* (Roy et al., 2025) represents the intermediate fusion baseline. For each configuration, median coverage probabilities are computed using completely held-out test data and summarized over 25 replications. Coracle maintains high and stable coverage across early, intermediate, and late fusion architectures, whereas non-conformal fusion methods exhibit noticeable deviations from nominal coverage.

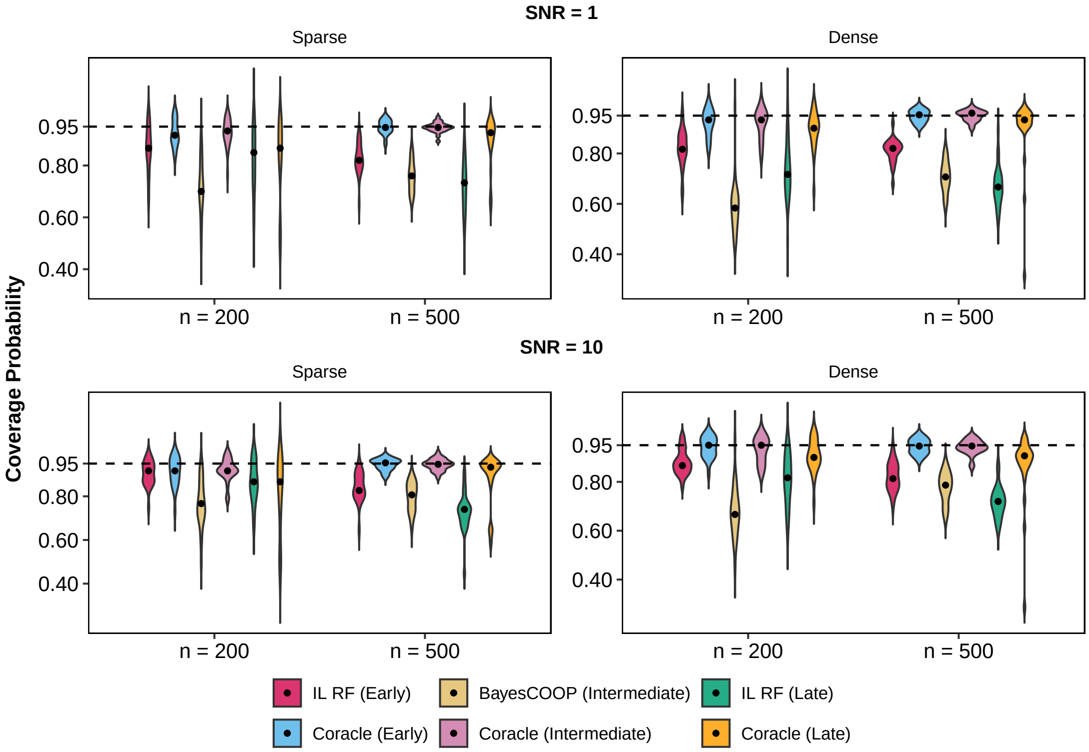


Figure S2: **Data-driven, non-factor model–based simulations with random forest as base learner for late fusion in high- and low-SNR settings**. Coverage performance for Coracle and published non-conformal baselines is evaluated under two signal-to-noise ratios (SNR = 1 and 10), two sample sizes ($n = 200, 500$), and both sparse and dense coefficient regimes. Here, *IL RF* denotes *IntegratedLearner* (Mallick et al., 2024) with random forest as the base learner (Breiman, 2001; Liaw and Wiener, 2002), and *BayesCOOP* (Roy et al., 2025) represents the intermediate fusion baseline. For each configuration, median coverage probabilities are computed using completely held-out test data and summarized over 25 replications. Coracle maintains high and stable coverage across early, intermediate, and late fusion architectures, whereas non-conformal fusion methods exhibit noticeable deviations from nominal coverage.

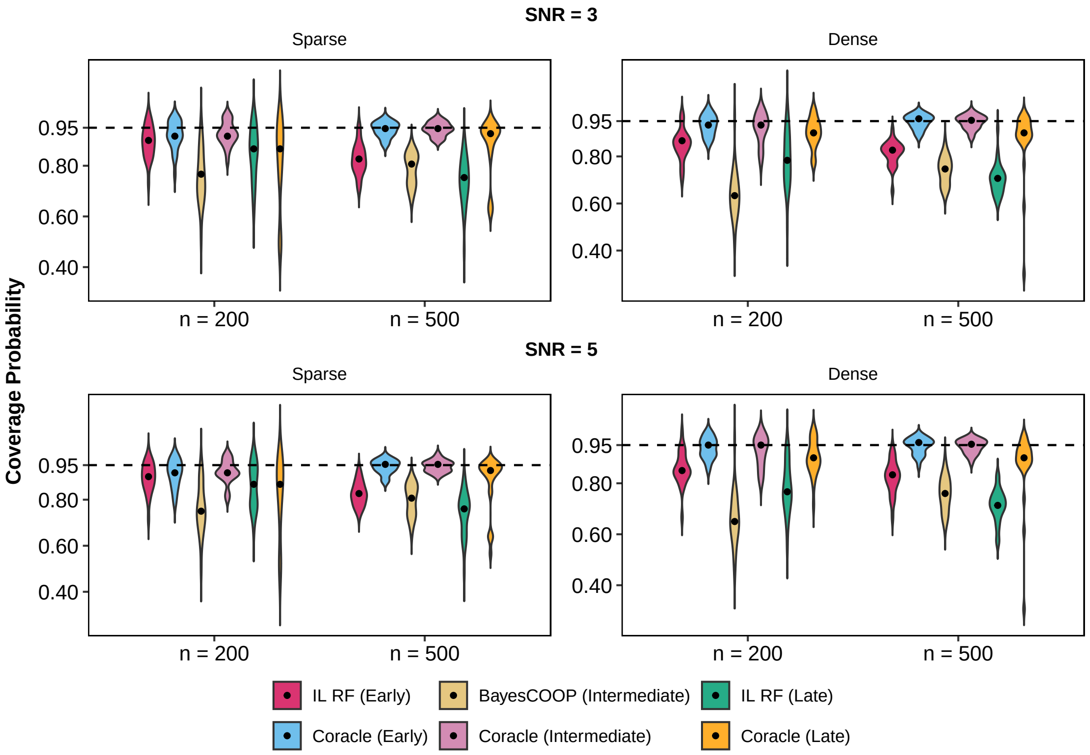


Figure S3: **Data-driven, non-factor model–based simulations with random forest as base learner for late fusion in moderate SNR settings**. Coverage performance for Coracle and published non-conformal baselines is evaluated under two signal-to-noise ratios (SNR $=$ 3 and 5), two sample sizes ($n = 200, 500$), and both sparse and dense coefficient regimes. Here, *IL RF* denotes *IntegratedLearner* (Mallick et al., 2024) with random forest as the base learner (Breiman, 2001; Liaw and Wiener, 2002), and *BayesCOOP* (Roy et al., 2025) represents the intermediate fusion baseline. For each configuration, median coverage probabilities are computed using completely held-out test data and summarized over 25 replications. Coracle maintains high and stable coverage across early, intermediate, and late fusion architectures, whereas non-conformal fusion methods exhibit noticeable deviations from nominal coverage.

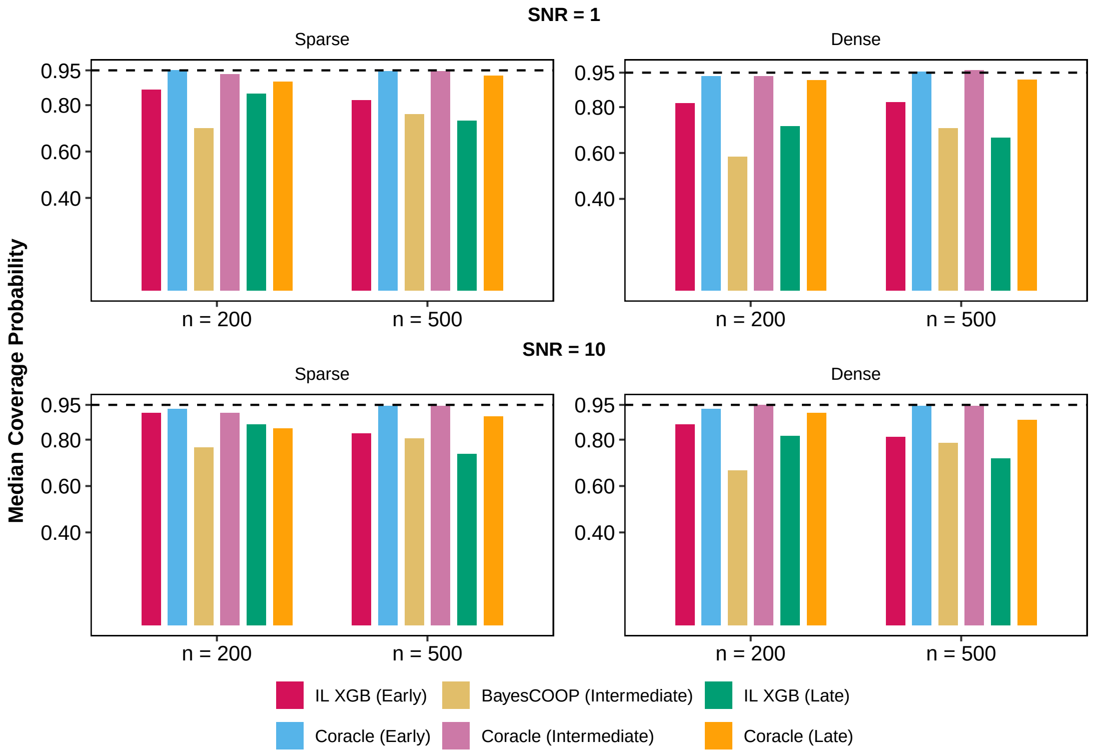


Figure S4: **Data-driven, non-factor model–based simulations with XGBoost as base learner for late fusion in high- and low-SNR settings.** Coverage performance for Coracle and published non-conformal baselines is evaluated under two signal-to-noise ratios (SNR = 1 and 10), two sample sizes ($n = 200, 500$), and both sparse and dense coefficient regimes. Here, *IL XGB* denotes *IntegratedLearner* (Mallick et al., 2024) with XGBoost as the base learner (Chen and Guestrin, 2016), and *BayesCOOP* (Roy et al., 2025) represents the intermediate fusion baseline. For each configuration, median coverage probabilities are computed using completely held-out test data and summarized over 25 replications. Coracle maintains stable near-nominal coverage across fusion strategies, while non-conformal methods remain noticeably miscalibrated.

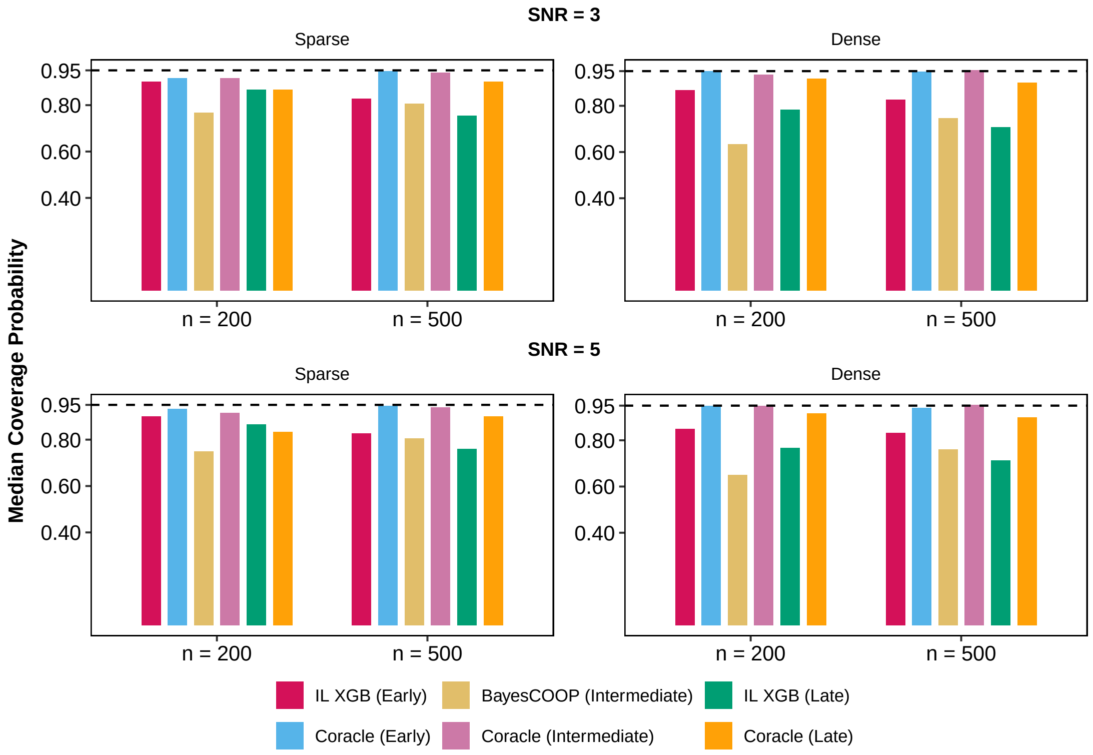


Figure S5: **Data-driven, non-factor model–based simulations with XGBoost as base learner for late fusion in moderate SNR settings.** Coverage performance for Coracle and published non-conformal baselines is evaluated under two signal-to-noise ratios (SNR $=$ 3 and 5), two sample sizes ($n = 200, 500$), and both sparse and dense coefficient regimes. Here, *IL XGB* denotes *IntegratedLearner* (Mallick et al., 2024) with XGBoost as the base learner (Chen and Guestrin, 2016), and *BayesCOOP* (Roy et al., 2025) represents the intermediate fusion baseline. For each configuration, median coverage probabilities are computed using completely held-out test data and summarized over 25 replications. Coracle maintains stable near-nominal coverage across fusion strategies, while non-conformal methods remain noticeably miscalibrated.

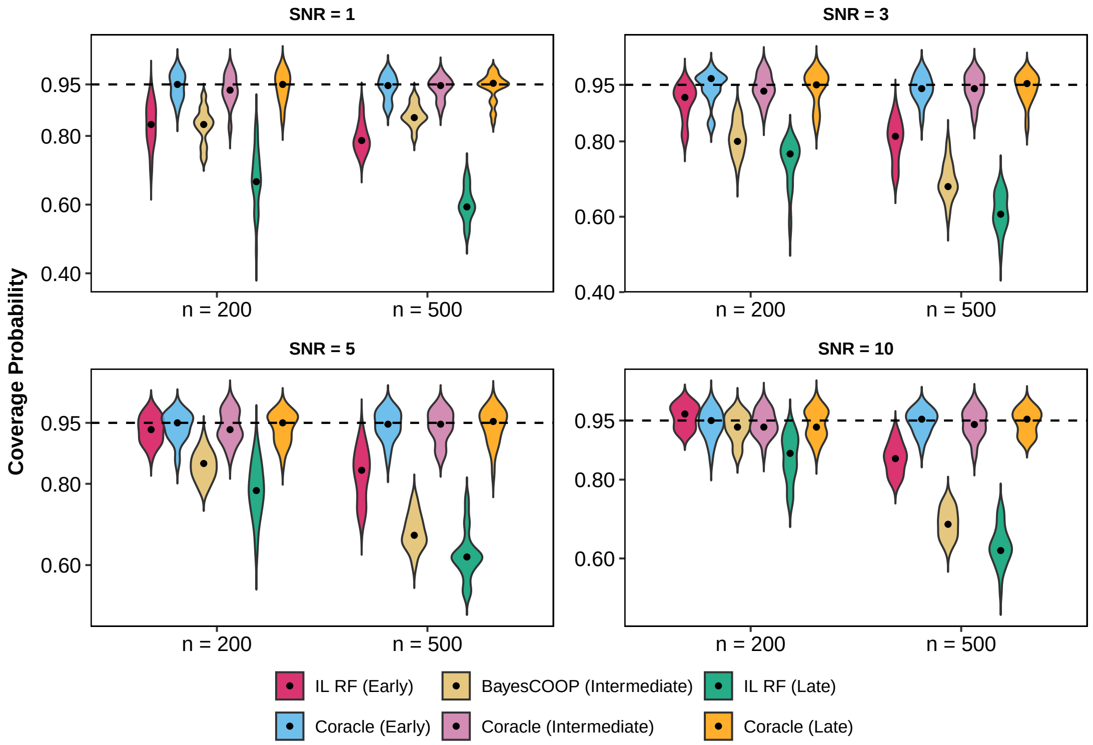


Figure S6: **Factor model–based simulations with random forest as base learner for late fusion.** Coverage performance for Coracle and published non-conformal baselines is evaluated under four signal-to-noise ratios (SNR = 1, 3, 5, 10) and two sample sizes ($n = 200, 500$). Here, *IL RF* denotes *IntegratedLearner* (Mallick et al., 2024) with random forest (Breiman, 2001; Liaw and Wiener, 2002) as the base learner, and *BayesCOOP* (Roy et al., 2025) represents the intermediate fusion baseline. For each configuration, median coverage probabilities are computed using completely held-out test data and summarized over 25 replications. Coracle consistently achieves near-nominal coverage across all fusion architectures, whereas non-conformal alternatives exhibit substantial miscalibration.

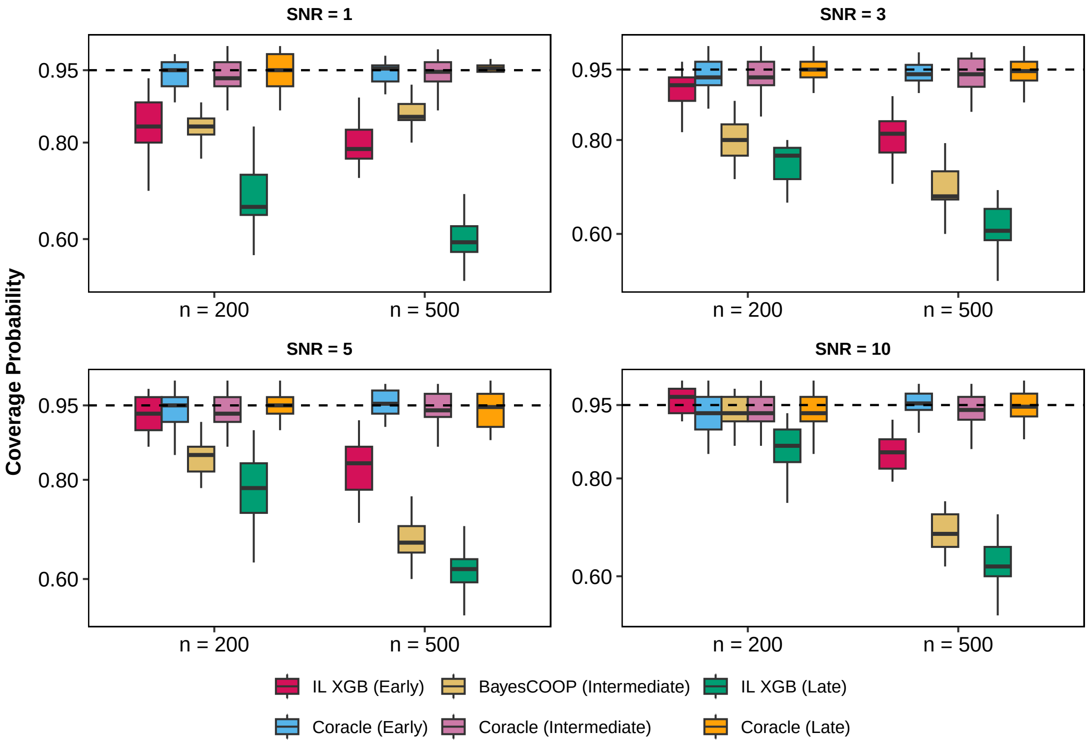


Figure S7: **Factor model–based simulations with XGBoost as base learner for late fusion.** Coverage performance for Coracle and published non-conformal baselines is evaluated under four signal-to-noise ratios (SNR = 1, 3, 5, 10) and two sample sizes ($n = 200, 500$). Here, *IL XGB* denotes *IntegratedLearner* (Mallick et al., 2024) with XGBoost (Chen and Guestrin, 2016) as the base learner, and *BayesCOOP* (Roy et al., 2025) represents the intermediate fusion baseline. For each configuration, median coverage probabilities are computed using completely held-out test data and summarized over 25 replications. Coracle consistently achieves near-nominal coverage across all fusion architectures, whereas non-conformal alternatives exhibit substantial miscalibration.

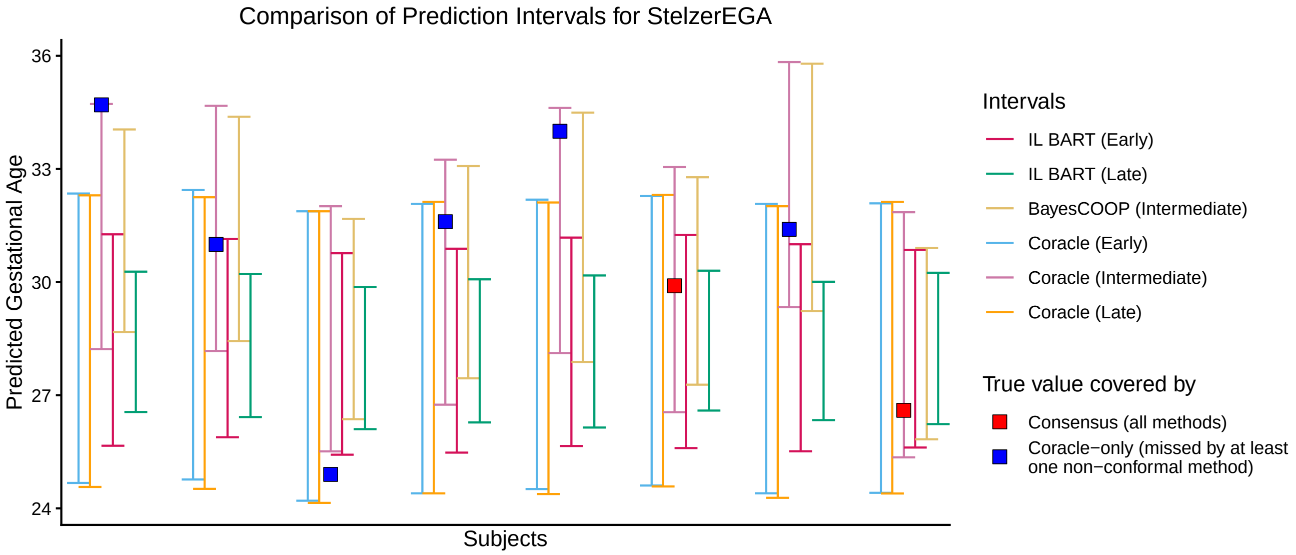


Figure S8: **Prediction interval comparison for the StelzerEGA dataset (Stelzer et al., 2021).** Subject-level prediction intervals for gestational age are shown for representative early, intermediate, and late fusion base models and their corresponding Coracle-conformalized versions, evaluated on completely independent validation data. Here, *IL BART* denotes *IntegratedLearner* (Mallick et al., 2024) with BART (Chipman et al., 2010) as the base learner (shown under early and late fusion), and *BayesCOOP* (Roy et al., 2025) denotes the Bayesian cooperative learning model used as the intermediate fusion baseline. For each subject, the true value is indicated, along with markers distinguishing cases in which all methods cover the true value from those in which Coracle achieves valid coverage but at least one non-conformal approach fails to do so.
